# Induction and antiviral activity of human β-defensin 3 against adenovirus and influenza A virus *in vitro* infection of human airway epithelial cells

**DOI:** 10.64898/2025.12.19.695592

**Authors:** Miguel Ángel Galván-Morales, Julio Santiago, Gerardo Corzo, Ligia Luz Corrales-García, Luis Manuel Teran

**Author notes:** Corresponding author, (w. city code) or 52 6465 4645. https://www.researchgate.net/profile/Julio-Santiago-8. https://www.researchgate.net/profile/Gerardo-Corzo, 5266.

## Abstract

**Background:** Respiratory pathogens such as adenoviruses (AdV) and influenza A viruses (IAV) can cause serious upper and lower respiratory infections in infants, the elderly, and immunocompromised individuals. However, options for antiviral drugs targeting respiratory viruses are limited. Human beta-defensins (hBDs) are disulfide-rich peptides that demonstrate broad antimicrobial activity against bacteria, fungi, and some viruses, while also playing immunomodulatory roles. Among defensins, human beta-defensins (hBDs) are the most prevalent, with six peptides identified: hBD-1 through hBD-6. Nonetheless, the production of hBD-3 during respiratory virus infections has not been extensively studied, nor has the effect of hBD-3 on these infections.

**Objective:** This study investigated whether infection with AdV-5 or IAV induces hBD-3 expression in human airway epithelial cells and whether a recombinant form of hBD-3 (rhBD-3) can inhibit *in vitro* infection by these viruses, one DNA virus and one RNA virus.

**Methods:** *In vitro* models of human airway epithelial cells (A549 and HEp-2) infected with AdV-5 and IAV were established. *In vitro* expression of hBD-3 mRNA was assessed by RT-PCR; hBD-3 protein was examined using immunofluorescence and Western blot. The inhibition of viral infection by hBD-3 was quantified using cytopathic effect and plaque reduction assays. Additionally, epithelial cells of animal origin, MDCK and MDBK, were similarly tested for rHBD-3 antiviral activity against IAV infection.

**Results:** IAV and AdV infections significantly increased hBD-3 mRNA levels in A549 and HEp-2 cells. Immunofluorescence and Western blotting confirmed the presence of hBD-3 protein on the surface of IAV-infected cells. Cells infected with AdV-5 produced less hBD-3 protein. Additionally, rhBD-3 inhibited both AdV-5 and IAV infections in both cell lines at concentrations of 7.5-50 μg/mL, with the strongest effect at the highest concentration. Lastly, rhBD-3 also inhibited IAV infection in MDBK and MDCK cells within the same range.

**Conclusions:** *In vitro* infection of human airway epithelial cells with IAV and AdV induces hBD-3 mRNA and protein expression. In turn, rhBD-3 inhibits viral infection in a dose-dependent manner in HEp-2 and A549 cells against these viruses, an RNA virus and a DNA virus. hBD-3 is expressed and localized on the surface of infected cells. rHBD-3 also inhibits IAV infection in bovine and canine epithelial cells. Our findings suggest that hBD-3 plays a broad-spectrum role in defense against respiratory viruses in humans and animals and modulates the innate immune responses.

**Perspective:** This work supports the use of hBD-3 as a natural, broad-spectrum antiviral agent, either alone or in combination, for treating viral infections, particularly emerging respiratory agents such as IAV and coronaviruses, for viral pandemic preparedness and response.

**Data Summary:** There are no supporting external data.

**Impact Statement:** This study demonstrates *in vitro* induction of human beta defensin type 3 (hBD-3) in human airway epithelial cells (A549 and HEp-2 cells) during viral infection by the influenza A virus (IAV) and the human adenovirus type 5 (AdV-5), and inhibition of the viral replication on these cells by the hBD-3 in a recombinant form; this defensin also showed viral inhibition on bovine and canine epithelial cells; further, these are viruses from two very different families; one is an enveloped RNA virus, and the other is a naked DNA virus. Both results are novel observations in the biomedical literature. Our results suggest that this defensin plays a principal role in innate defense during natural respiratory viral infections in humans, and support the use of hBD-3 alone or in combination as a broad-spectrum antiviral agent for the treatment of these infections, especially for emerging agents such as IAV and coronaviruses, for viral pandemic preparedness and response. In fact, few antiviral drugs are currently approved for treating respiratory virus infections, with only specific inhibitors of influenza and respiratory syncytial viruses.

## Introduction

Viral respiratory diseases are a leading cause of lower respiratory infections, which pose a significant health burden worldwide, mainly affecting children and the elderly. These infections are caused by viruses such as the influenza A virus (IAV), coronaviruses, human adenoviruses (AdV), rhinoviruses, and respiratory syncytial virus, among others [Bender *et al*., 2024; WHO, 2025]. IAV, an enveloped RNA virus, is highly pathogenic, with seasonal strains infecting 3-5 million people annually and causing symptoms ranging from flu-like illness to severe disease. Additionally, IAV could potentially become a pandemic, as the last one occurred in 2009. Vaccination against IAV protects against infection in up to 86% of cases, and efforts are ongoing to improve seasonal vaccines due to the high mutation rate of RNA viruses, which leads to minor changes (antigenic drift) and major changes (antigenic shift). However, antiviral drugs for respiratory viruses are limited. Typically, only two antiviral agents are available for IAV: oseltamivir and amantadine (Jain *et al*., 2025). AdV (naked DNA viruses) not only causes lower respiratory tract disease but can also lead to gastrointestinal, urinary, meningoencephalitis, conjunctivitis, and cystitis. AdV infections spread easily among human populations, with outbreaks often occurring in crowded settings such as healthcare facilities, military bases, and schools. Currently, no vaccine is available for the general public, and no specific treatments have been developed for AdV (Crenshaw *et al*., 2019). Therefore, there is an urgent need for antivirals targeting IAV, AdV, and other respiratory viruses.

Defensins are small, cationic peptides with a molecular weight under 20 kDa that contain multiple cysteine residues. They play a role in modulating the innate immune response and defending against bacteria, fungi, and viruses. Additionally, they have immunomodulatory effects. These antimicrobial peptides were first identified in the 1980s by Lehrer and colleagues in mammalian neutrophil phagocytes. They have six conserved cysteine residues that form disulfide bonds, inducing conformational changes. Based on these structural characteristics, defensins are divided into three subfamilies: alpha (α-), beta (β-), and theta (θ) defensins. In humans, only α- and β-defensins are known to be present [Patterson-Delafield *et al*., 1980; Xu and Lu, 2020].

Human β-defensins (hBDs) are the most widely distributed defensins, with six peptides (hBD-1 to hBD-6) encoded by more than 30 genes clustered on chromosome 8p23.1 [Sun *et al*., 2006]. β-Defensin expression is transcriptionally regulated and limited to keratinocytes in the skin, as well as epithelial cells, leukocytes, glands, and Paneth cells. For example, hBD-3 is found in Langerhans cells and the skin, and the production of hBD-2-4 and hBD-3 is mediated by cytokines such as TNF-α and IFN-γ, respectively [Winter and Mathias, 2012; Hazlett and Wu, 2011; De Paula and Valente, 2018; Ding *et al*., 2009].

The mechanism of action of hBDs is multifunctional; for instance, in CD4+ T cells and macrophages, hBDs inhibit protein kinase C (PKC) activity. In HIV infections, hBDs interfere with nuclear import and virus transcription [Weinberg *et al*., 2006]. Additionally, by binding to the C1q peptide, hBD-1 and −2 inhibit the classical complement pathway (Groeneveld *et al*., 2007). In several models, such as HIV and AdV, hBDs directly interfere with viral infection by binding to viral receptor glycans or adherence proteins on the viral surface (Gong *et al*., 2010). It has also been shown to damage cellular membranes by binding cations and inhibiting both endogenous and exogenous potassium channels [Zasloff, 2002; Zhang *et al*., 2018; Li *et al*., 2024].

hBD-3 disrupts the biosynthesis of bacterial cell walls in Gram-negative and Gram-positive bacteria by binding to lipid-II-rich regions of the cell wall [Sass *et al*., 2010]. A similar mechanism has been demonstrated for the fungal defensin plectasin [Schneider *et al*., 2010]. hBD-3 inhibits human immunodeficiency virus type 1 (HIV-1) replication by downregulating the HIV-1 coreceptor CXCR4 (Quiñones-Mateu *et al*., 2003). Additionally, hBD-3 interferes with viral fusion processes mediated by the IAV hemagglutinin (HA), the penton fiber in AdV infection, the baculovirus gp64, and the Sindbis virus E1 protein (Leikina *et al*., 2005).

Recently, in mouse models and Madin-Darby canine kidney (MDCK) cells, the recombinant mouse BD-3 (rmBD) was tested at various stages of IAV A/PR/8/34 (H1N1) infection to evaluate its potential properties [Jiang *et al*., 2012; Skalickova *et al*., 2015]. In 2015, HeLa cells and mice infected with Coxsackie virus CVB3 treated with rMBD3 showed protective effects against viral infection *in vitro* and myocarditis in mice [Jiang *et al*., 2015].

After a thorough review, we found that neither hBD-3 nor its recombinant homolog has been tested for antiviral activity against IAV or AdV in human airway cell models. Therefore, we decided to study the antiviral activity of hBD-3 during *in vitro* infection with these viruses. First, we investigate whether infections with AdV type 5 (AdV-5) and IAV induce hBD-3 in human airway epithelial cells (HEp-2 and A549), and then whether rhBD-3 blocks IAV and AdV infection: an enveloped RNA virus and a naked DNA virus.

## Materials and methods

### Expression and purification of recombinant hBD-3

The expression and purification of recombinant hBD-3 (rhBD-3) followed our previous work [Corrales *et al*., 2013], with some modifications: the recombinant defensin was vacuum dried, and 500 µg of cystine-reduced recombinant defensin was resuspended in 2 mL of 6 M GndHCl (pH 2). Then, 1.8 and 0.36 mg of reduced glutathione (GSH) and oxidized glutathione (GSSG), respectively, were added. The reduced defensin solution was further diluted to 6 mL with 3 mL of distilled water and 1 mL of 0.3 M Tris buffer, pH 8.0. Next, the recombinant product was allowed to fold for 24 h at room temperature in a folding solution containing 2 M GndHCl in 0.05 M Tris buffer, pH 8.0, with 1 mM GSH and 0.1 mM GSSG. Recombinant hBD-3 was then purified by HPLC using a C18 column (Vydac C18, 5 µm, 300 Å, 4.6 x 250 mm) at a flow rate of 1 mL/min. The molecular weight of rhBD-3 was confirmed by mass spectrometry using an LCQ Fleet Ion Trap (Thermo Scientific, Barrington, IL, USA). Finally, recombinant hBD-3 was freeze-dried, stored at −20 °C, and reserved for subsequent analysis. The peptide concentration was determined by measuring absorbance at 280 nm using its respective molar extinction coefficient.

### Cell lines

The cell lines used for the assays included A549 human alveolar epithelial cells (ATCC, CCL-185), HEp-2 human laryngeal epithelial carcinoma cells (ATCC, CCL-23), MDBK cells from Madin-Darby Bovine Kidney (ATCC, CCL-22), and MDCK cells from Madin-Darby Canine Kidney (NBL-2) (ATCC, CCL-34). Cells were cultured in 25 cm2 flasks (Corning Life Sciences, Catalog Number: 431463) containing Modified Eagle’s Medium (MEM) (Gibco Cat.12492013) supplemented with 10% fetal bovine serum (Thermo Fisher Scientific, Cat. 16000069), along with antibiotics: penicillin 100 U/mL, streptomycin 100 μg/mL, and amphotericin-B 1 μg/mL (Sigma-Aldrich, Cat: A5955-100mL). Cultures were incubated at 37 °C with 5.0% CO_2_ (v/v) in humidity-saturated air; all subsequent 37 °C incubations were conducted under these conditions. Cells were transferred to 24- and 96-well plates for experimental procedures, including viral titration, Western blot, RT-PCR, immunofluorescence, and plaque assays, to assess the effects of rhBD-3. Cell cultures, fetal bovine serum, and media were regularly checked for bacterial contamination via standard bacterial culture methods and for mycoplasma contamination using the EZ-PCR Mycoplasma Detection Kit (SKU: 20-700-20, Sartorius, Biopharma, Lab, Applied & Life Sciences).

### IAV and AdV propagation and titration

Influenza virus type A/WSN/33 (H1N1) (VR-219™) and AdV-5 (VR-1516™) were obtained from ATCC®. Each virus was propagated in monolayers of HEp-2, A549, MDBK and MDCK cell lines at 90% confluence in 25 cm² flasks. The monolayers were washed twice with sterile phosphate-buffered saline (PBS), and then inoculated at a multiplicity of infection (MOI) of 0.01 virus per cell for IAV or with 10 tissue culture infectious doses 50% (TCID_50_) per flask for AdV-5. The monolayers were incubated for one h at 37 °C. Inocula were then discarded, and fresh medium was added to the flasks. For IAV, 0.050 mL of trypsin (0.05%, Gibco, 11580626) was added to the culture medium; in all experiments with IAV, trypsin 0.05% was used at the same concentration: 50 Ll/5 mL of MEM. All cultures were incubated at 37 °C until cytopathic effect (CPE) was observed. Infected cell suspensions were centrifuged at 750 x *g* for 10 min at 4°C. The supernatant was stored in 1.5 mL aliquots with 10% DMSO (D8418, Sigma-Aldrich) at −70 °C for cryopreservation.

For plaque assays, serial 1:10 dilutions of both viruses were prepared in MEM medium. Confluent monolayers of cell lines were seeded in 24-well plates, inoculated with 100 µL of each dilution in triplicate, and incubated for one h at 37 °C. After incubation, supernatants were discarded, monolayers were washed with sterile PBS, and 1 mL of 1.5% methylcellulose (MC) in MEM without FBS was added to each well. MC viscosity prevents viruses from spreading across the well, ensuring they propagate only by contact and form well-defined plaques. Cells were incubated until lytic plaques appeared, typically between 48 and 72 h. MC medium was removed, and the cells were gently washed with PBS. Then they were incubated with 200 μL of 75% methanol for 1 min, followed by a 15-min treatment with 1% crystal violet (C0775, Sigma). After washing with water, plaques were counted under a microscope (Leica, DM IL LED Inverted Microscope 090135.001) to quantify the plaque-forming units (pfu). Three replicates of each assay were performed for each experiment.

### RNA Extraction and cDNA Synthesis

mRNA extraction for reverse transcription. The expression of hBD-3 mRNA in HEp-2 and A549 cells following infection with IAV and AdV-5 was analyzed by reverse transcription-PCR (RT-PCR). Monolayers of HEp-2, A549, and MDCK cells at 90% confluence in 25 cm² flasks were infected with IAV or AdV-5 as described previously. The monolayers were washed with PBS and then inoculated with IAV at an MOI of 0.01 virus per cell or with AdV-5 at 10 TCID_50_ per flask. Cells were incubated for 1 h, then the inocula were discarded, and fresh medium was added to the flasks. For IAV, 0.050 mL of trypsin (0.05%) was added per flask. All cultures were incubated until cytopathic effect (CPE) was observed.

Total RNA from infected cells was extracted using the commercial SV total RNA isolation System (Promega, Madison, USA) according to the manufacturer’s instructions. cDNA synthesis was performed in a total volume of 20 μL using sterile microtubes of 200 μL, 5.0 μg/mL of total RNA, 500 μg/mL of Oligo d(T) (Invitrogen, CA, USA), 10 mM of dNTPs mix (Invitrogen, CA, USA), and DEPC-treated water to reach 20 μL. Denaturation was carried out at 65°C for 5 min. Samples were then rapidly centrifuged and incubated for 2 min at 42°C. Next, 1.0 μL (200 units) of SuperScript II enzyme (Invitrogen) was added, and the mixture was incubated for 50 min at 42°C. The reaction was completed by incubating for 15 min at 70°C.

RT-PCR. Total RNA was reverse transcribed using Moloney murine leukemia virus (M-MLV) reverse transcriptase (Invitrogen, Karlsruhe, Germany). RT-PCR amplified the resulting cDNA with Taq polymerase (Qiagen). Primers specific for GAPDH (glyceraldehyde-3-phosphate dehydrogenase; 5’-CCACCCATGGCAAATTCCATGGCA-3’ and 5’-TCTAGACGGGCAGGTCAGGTCCACC-3’) and hBD-3 (5’-TCTCAGCGTGGGGTGAAGC-3’ and 5’-CGGCCGCCTCTGACTCTG-3’) were used. These primers were obtained from TIB Molbiol (Berlin, Germany). After amplification, PCR products were separated by electrophoresis on 2% agarose gels, stained with ethidium bromide, and visualized (ChemiDoc XRS+ Gel Imaging System). GAPDH expression served as a control to verify equal amounts of cDNA were used in each experiment.

### Analysis of hBD-3 expression by Western blot

Confluent monolayers of HEp-2 and A549 cells in 96-well plates were infected with IAV and AdV-5 at infectious doses of 0.01 MOI and ten TCID_50_, respectively, for 48 h. Protein extraction was performed using 1X RIPA (Cell Signaling Technology, 9806) according to the manufacturer’s instructions. Proteins were quantified by the Bradford method. They were separated by SDS-PAGE on 18% polyacrylamide gels and transferred to PVDF membranes (BioRad, 1620177) using a Semi-Dry Transfer Cell (BioRad, Trans-Blot® SD) at 150 V for 1 h. The membranes were blocked with 5% nonfat milk in TPBS and incubated overnight at 4 °C with primary goat anti-hBD-3 (R&D Systems, AF4435) at a 1:1000 dilution. After washing, they were incubated with a secondary antibody, fluorescein isothiocyanate-labeled rabbit anti-goat IgG (Santa Cruz Biotechnology, sc-2356), at a 1:1000 dilution. The reaction was detected by chemiluminescence on X-ray film with Luminol (Santa Cruz Biotechnology, sc-2048). The bands were analyzed by densitometry using ImageJ (U.S. National Institutes of Health).

### Immunofluorescence of hBD-3 in the infected cell lines

After the viral infection described above, A549 and HEp-2 cells were scraped from culture flasks and placed onto glass slides. The cells were fixed with 1% acetone, air-dried, and stained with a primary goat anti-hBD-3 antibody (R&D Systems, AF4435) at a concentration of 2.5 µg/mL, followed by a secondary rabbit anti-goat IgG antibody conjugated with fluorescein isothiocyanate (Santa Cruz Biotechnology, sc-2356) at 1.5 µg/mL. Both antibodies were diluted in phosphate-buffered saline (PBS) containing 0.6% EDTA solution, and 5% BSA. After each antibody incubation, the slides were gently washed with PBS to remove any unbound antibody. Each slide was then counterstained with 50 µL of propidium iodide (Sigma-Aldrich, P4864). The slides were observed and images captured using a fluorescence microscope (Olympus BX60) equipped with an integrated Leica DC100 camera and the Leica Application Suite X software.

### rhBD-3 antiviral activity assay

To investigate the activity of recombinant human β-defensin 3 (rHBD-3) on *in vitro* infections caused by IAV and AdV-5, the A549 and HEp-2 cell lines were cultured in 96-well plates. The rhBD-3 was dissolved in PB (10 mM phosphate buffer; 8.2 mM Na_2_HPO_4_, 1.8 mM KH_2_PO_4_, pH 7.4) and diluted in MEM. In a first step, viruses were incubated with rhBD-3 for 60 min before infecting cells. For this, 25 µl of AdV-5 (10 TCID_50_ units per well) or IAV (200 pfu for 20,000 cells, MOI 0.01) were incubated with 25 µl of each hBD-3 dilution in serum-free culture medium (MEM) for 60 min on ice bath. The mixtures were then added to cell monolayers in 96-well plates in triplicate and incubated for one h at 37 °C. Inocula were discarded and, in the second step, infected cells were incubated for 48 h with medium containing the same concentration of hBD-3.

Concentrations of rhBD-3 were 7.5, 12,5, 25, and 50 μg/mL, within the range previously reported [Salvatore *et al*., 2007; Smith *et al*., 2010]. In parallel, wells were infected with dilutions of each virus alone to establish a kinetics curve for cell viability after infection. After 48 h, the culture medium was removed, and the cells were gently washed with PBS. Cells were stained with tetrazolium blue (Sigma, M2128, 5 mg/mL in PBS). The dark blue crystals formed were solubilized in acidified isopropanol, and the absorbance of the solution at 570 nm was measured using an ELISA reader. Data were expressed as the percentage of cell viability and analyzed under nonlinear regression (Excel software). Uninfected cell controls incubated with rhBD-3 were used to assess defensin cytotoxicity. MDCK and MDBK cell lines were used to replicate these assays.

For the plaque inhibition assay, MDBK and MDCK epithelial cell lines were cultured in 24-well plates, and experiments were conducted as described above. A consistent volume of 100 μL of IAV (100 pfu) was added to the monolayers for infection. Cells were incubated with rHBD-3 as previously described, and plaques were counted following the procedure outlined in the viral propagation and titration section.

### Statistical analysis

Results are presented as the mean with the standard error (SE) from three replicates. The data were analyzed using a T-test to assess significant differences among group means. Graphs were created using RStudio software.

## Results

### Influenza A virus and adenovirus induce hBD-3 mRNA in HEp-2 and A549 cells

We hypothesized that AdV and IAV infections might induce hBD-3 synthesis in human epithelial cell lines. First, we examined whether IAV and AdV trigger hBD-3 mRNA expression in A549 and HEp-2 cells. RT-PCR analysis revealed that hBD-3 mRNA was significantly increased in both cell lines after infection with IAV, showing two strong bands on the gel, while AdV-5 induced less hBD-3 mRNA, producing two faint bands compared to the IAV infection (Fig. 1A and B); relative to AdV, IAV-induced hBD-3 mRNA expression was 5-fold higher in A549 cells and 3-fold higher in HEp-2 cells. Uninfected A549 and HEp-2 cells exhibited minimal constitutive hBD-3 mRNA expression.

**Figure 1.**
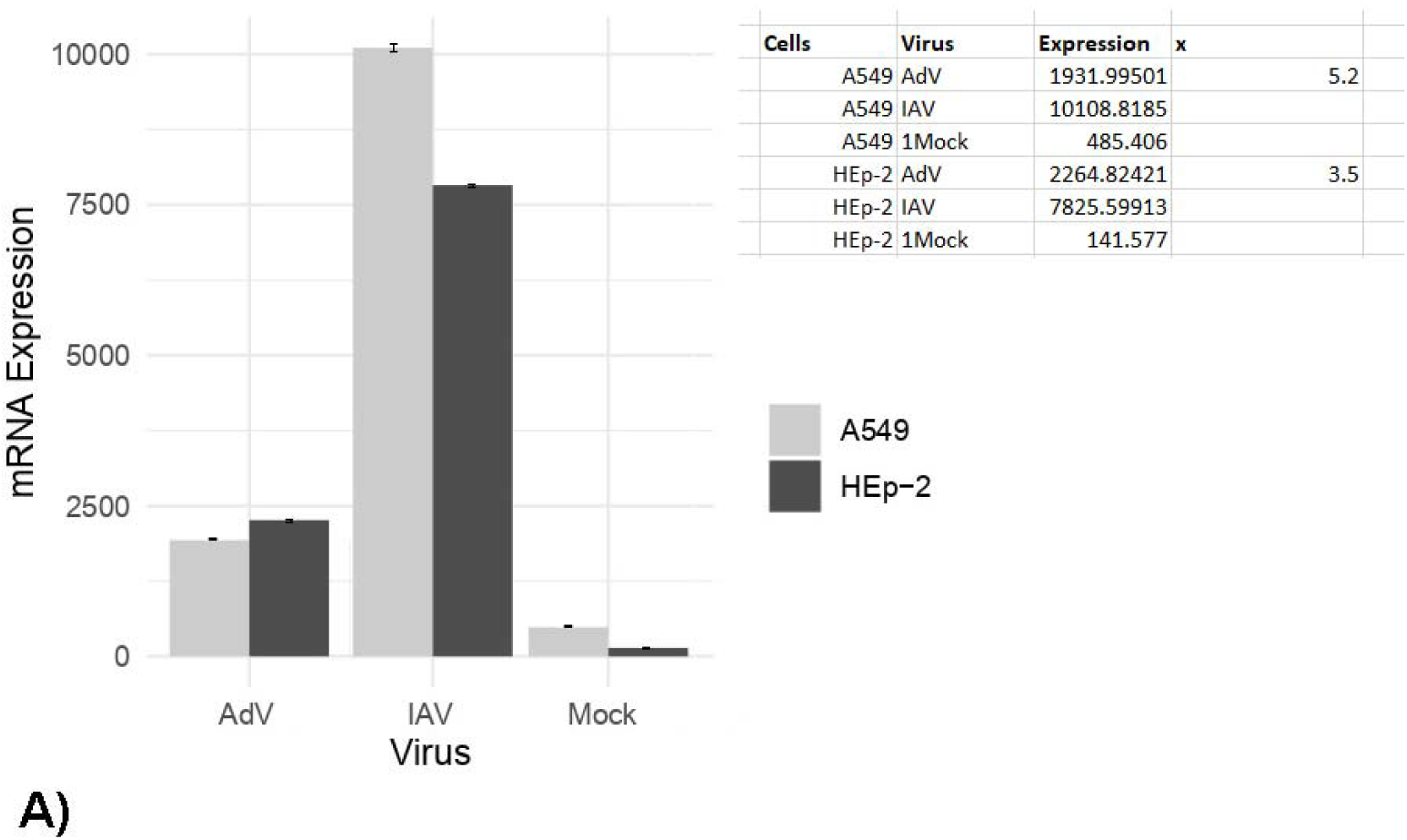
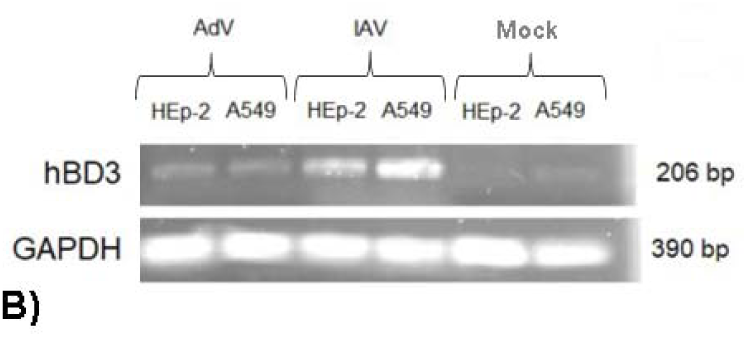
hBD-3 mRNA expression in HEp-2 and A549 cells after 48 hours of infection with IAV and AdV-5. Semi-quantitative RT-PCR products of hBD-3 following infection with IAV at an MOI of 0.01 (pfu/cell) and AdV-5 at 10 TCID_50_ per flask. Band intensities were measured by densitometry on gel images using the ImageJ software. A) Bars 1-2: A549 and HEp-2 cells infected with AdV; Bars 3-4: cell lines infected with IAV; Bars 5-6: mock-infected lines. Data are presented as means ± SE (p < 0.004). Results were quantified, and significant differences were compared quantitatively, as shown in the top box. B) Agarose gel electrophoresis of the respective hBD-3 mRNA and GAPDH amplimers. GAPDH was used as a constitutive control to normalize cDNA levels.

### Protein expression of hBD-3 in infected cell lines

Next, after confirming mRNA expression, we analyzed hBD-3 protein levels using Western blot. Similarly, A549 and HEp-2 cell lines infected with AdV-5 and IAV were examined by one-dimensional gel electrophoresis with specific antibodies. hBD-3 positive bands from HEp-2 and A549 cells infected with influenza A virus were observed; however, no positive bands were detected for adenovirus infection in either cell type (Fig. 2A and B). Uninfected cell lines showed no bands.

**Figure 2.**
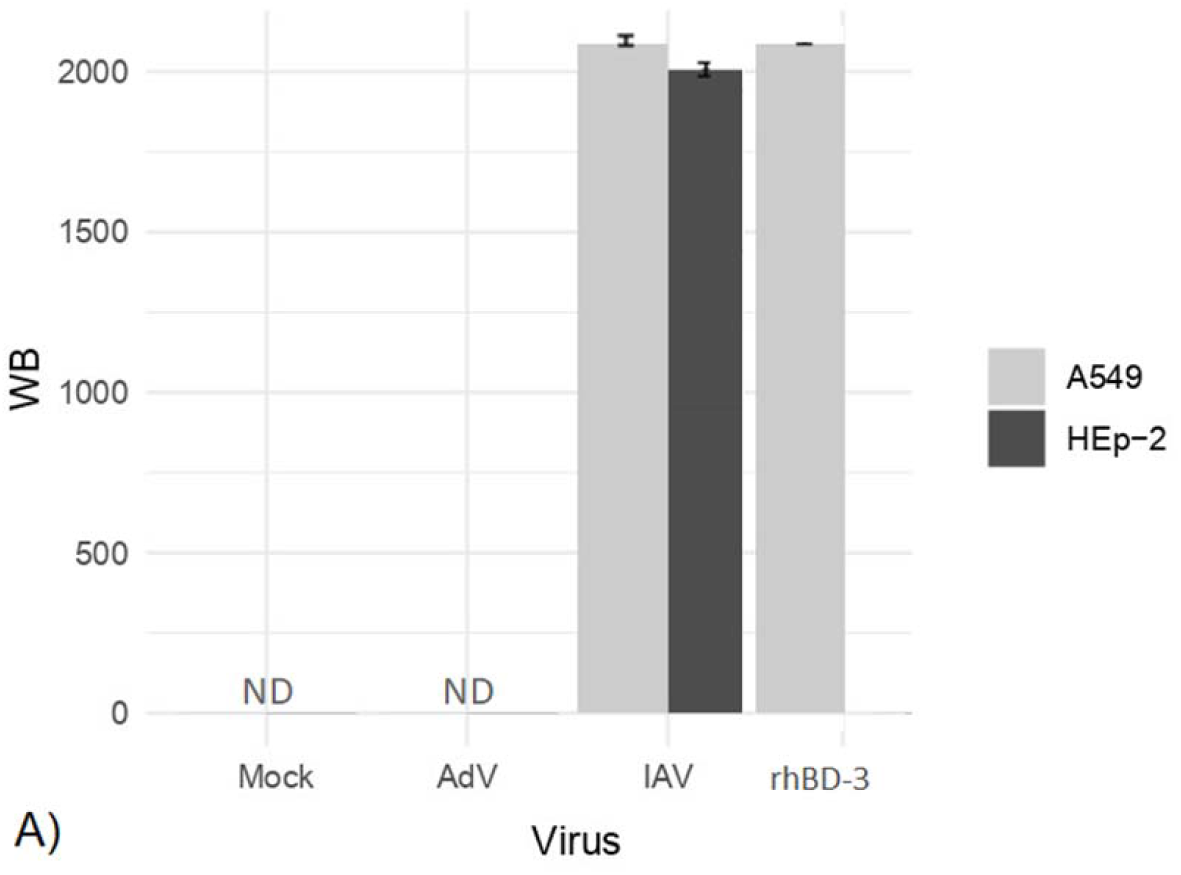
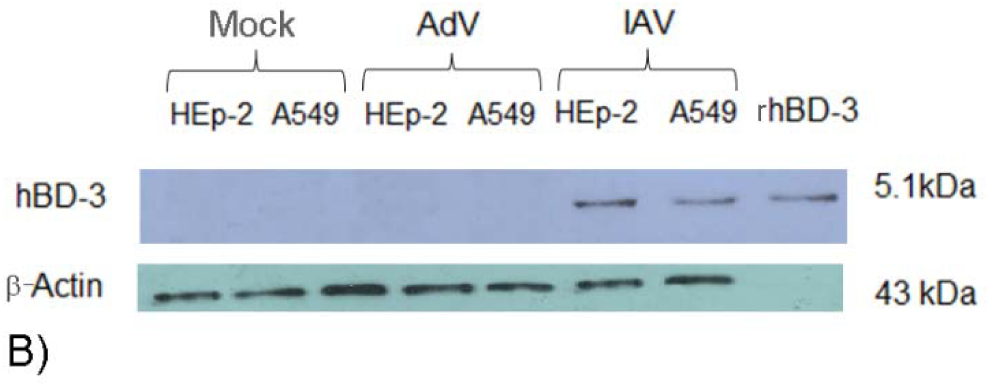
hBD-3 protein expression in A549 and HEp-2 cells infected with AdV-5 or IAV 48 hours post-infection. (A) Densitometry analysis of protein levels: the first two bars represent mock-infected cells; the third and fourth bars show adenovirus-infected cells (with undetectable hBD-3 protein, ND); the next two bars depict IAV-infected cells. Data from three independent experiments are presented as mean ± SE (p < 0.002 between IAV values and other data). (B) hBD-3 immunoblot of cells infected with AdV and IAV; lanes 5-6 display clear expression of hBD-3 in IAV-infected cells. Additionally, a recombinant hBD-3 protein (rhBD-3) control was loaded in the last lane.

### Immunofluorescence of hBD-3 in HEp-2 and A549 cell lines

Next, we examined the cellular localization of hBD-3 induced by viral infection. Immunofluorescence specific to hBD-3 was performed on HEp-2 and A549 cells infected with IAV and AdV-5. Uninfected cell lines did not show distinguishable fluorescence signals (fig. 3a and b). HEp-2 and A549 cells infected with IAV showed evident green fluorescence signals emitted by hBD-3 localized on the surface of infected cells (Figs. 3c and d). Fluorescent signals were partially present on AdV-infected cells (Fig. 3e and f). Green fluorescence signals were also localized on the surface of rhBD-3-treated cell lines, similar to infected cells (3g-h). Zoom-in images of IAV-infected cells were taken to confirm the location of hBD-3 with greater resolution (Fig. 3i-l).

**Figure 3.**
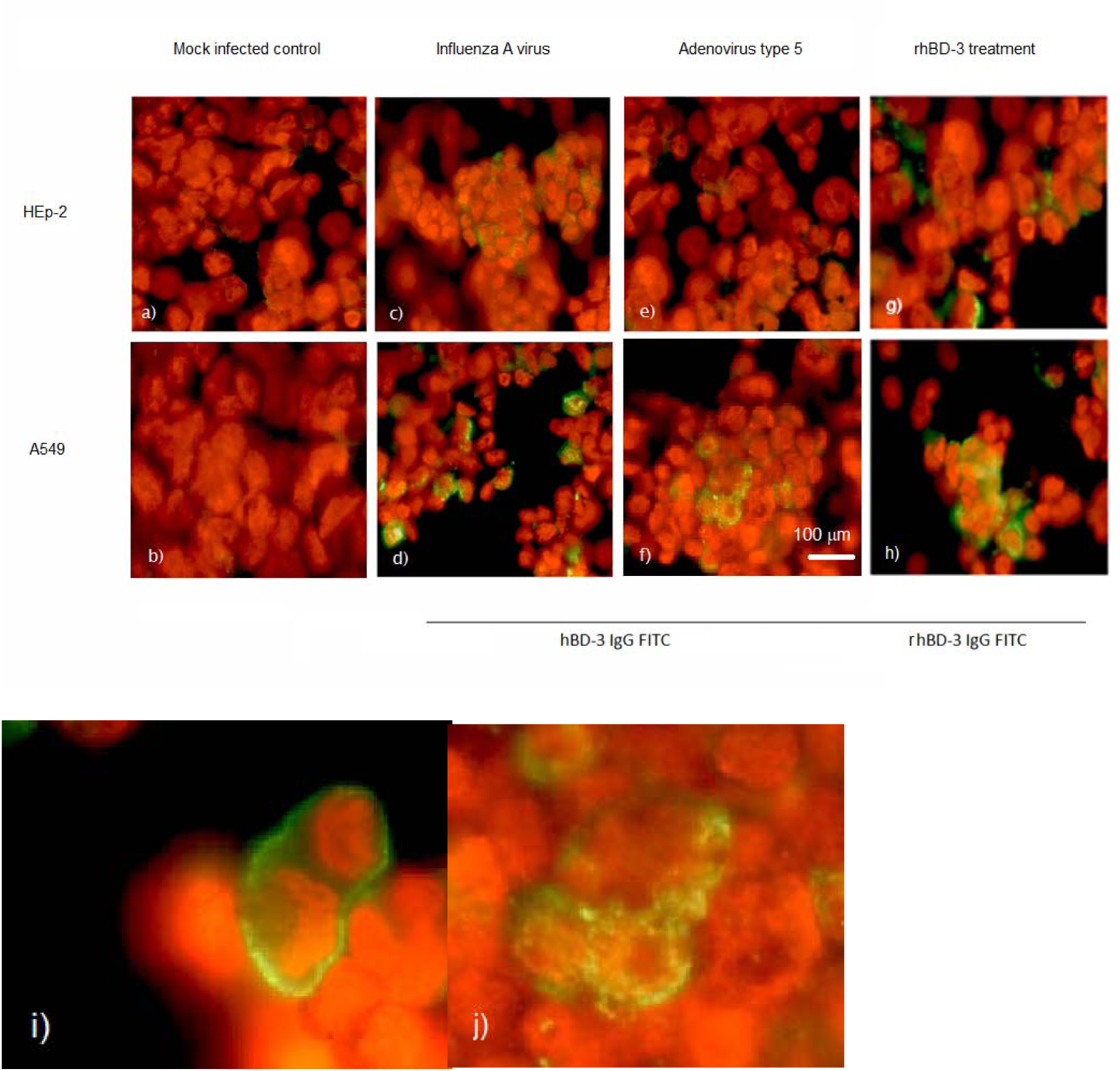
hBD-3 immunofluorescence in HEp-2 and A549 cells. a) Uninfected HEp-2 cells, and b) A549 cells stained with conjugated antibody and counterstained with propidium iodide to visualize whole cells. c and d) HEp-2 and A549 cells infected with IAV. e and f) Cell lines infected with AdV-5. g and h) Uninfected cells treated with rhBD-3 protein (positive control). Cells were stained with a specific anti-hBD-3 antibody conjugated with FITC and examined in an epifluorescence microscope. Scale bar = 100 μm (in f). i) and j) Close-ups of IAV-infected A549 cells showing the location of hBD-3.

### Antiviral activity of rhBD-3

To determine whether human airway epithelial cells could be protected from viral respiratory infections, we treated A549 and HEp-2 cells with a recombinant form of hBD-3 (rhBD-3) before infecting them with IAV and AdV. Cells treated with rhBD-3 were protected against AdV infection across a range of 7.5-50 μg/mL. Figures 4a and 4b show increasing viability levels in A549 and HEp-2 cells infected with AdV, which directly correlated with defensin concentration, with the highest effect at 50 μg/mL, indicating a dose-dependent inhibitory effect with a nonlinear correlation that tends to plateau at higher concentrations. Similar results were observed with IAV (Figs. 4c and 4d). Fig. 4e shows the antiviral effect of rhBD-3 on AdV infection of A549 cells, which appears as a gradual reduction in cell lysis visible without magnification. Similar effects are seen in Fig. 4f) with AdV infection in HEp-2 cells, Fig. 4g) with IAV infection in A549 cells, and Fig. 4h) with IAV infection in HEp-2 cells.

**Figure 4.**
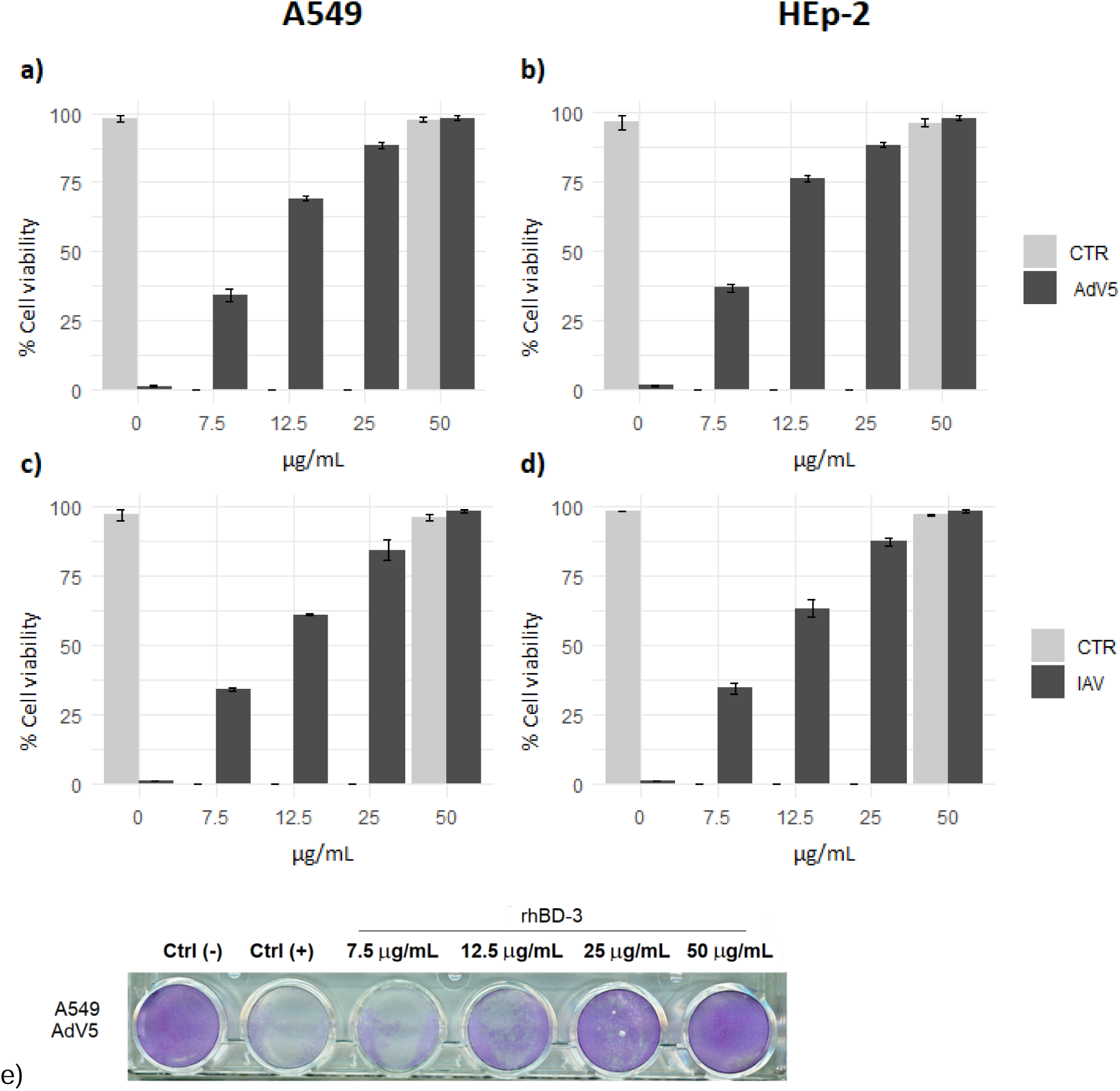
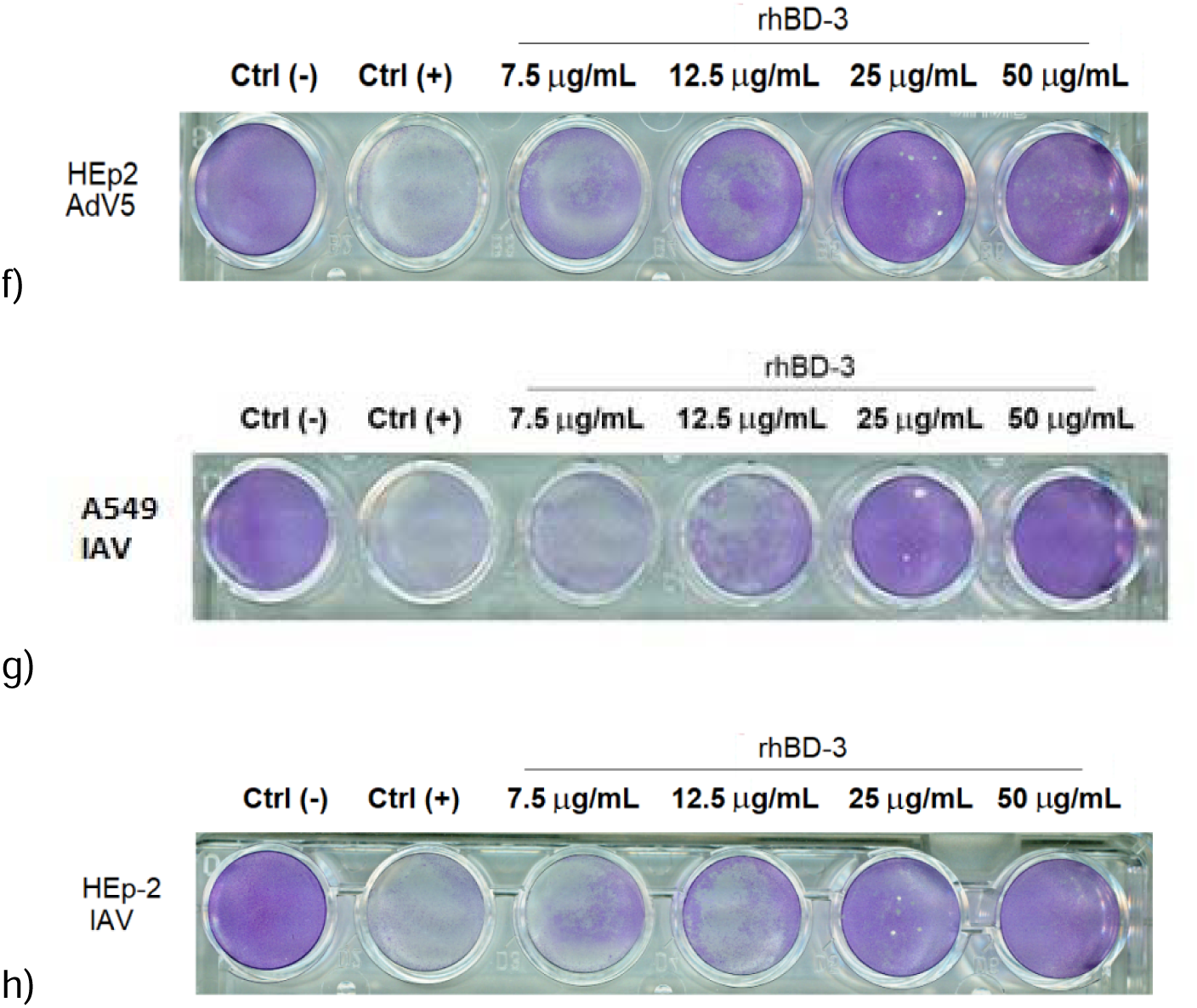
*In vitro* antiviral activity of hBD-3 against influenza A virus (IAV) and adenovirus type 5 (AdV-5). (a) Cell viability of A549 cells treated with rhBD-3 at 7.5, 12,5, 25, and 50 μg/mL, and infected with AdV-5. rhBD-3 inhibited AdV infection in A549 cells in a dose-dependent manner, in a nonlinear correlation. (b) Defensin also inhibited AdV in HEp-2 cells. (c) Cell viability of A549 cells and (d) HEp-2 cells treated with hBD-3 and infected with IAV. IAV infection was also inhibited in both cell lines. Data correspond to three individual experiments expressed as means ± SE. e) y f) Monolayers of A549 and HEp-2 cells treated with hBD-3 as indicated and infected with AdV-5. In the first well, the control cells are not infected; in the second through sixth wells, distinct concentrations of rhBD-3 are indicated. Gradual inhibition of the cytopathic effect is visually evident in the sequential cultures. g) and h) Monolayers of A549 and HEp-2 cells treated with defensin and infected with IAV with similar inhibitions for the viral infection.

To study the effect of hBD-3 on animal cells, MDCK and MDBK cells were also infected with IAV under similar hBD-3 concentrations. Consistently, rhBD-3 also effectively blocked IAV infection in these animal cell lines, with inhibitory curves closely resembling those observed in human cell lines. Figures 5a and b show the dose-dependent inhibitory effects on IAV and AdV-5 respectively, from 7.5 to 50 mg/mL doses. A concentration of 50 mg/mL inhibits almost the 100 percent of the pfu of IAV. Figure 5c and d show the viral cultures on cells treated with the rhBD-3, and the effects are visible without magnification.

**Figure 5.**
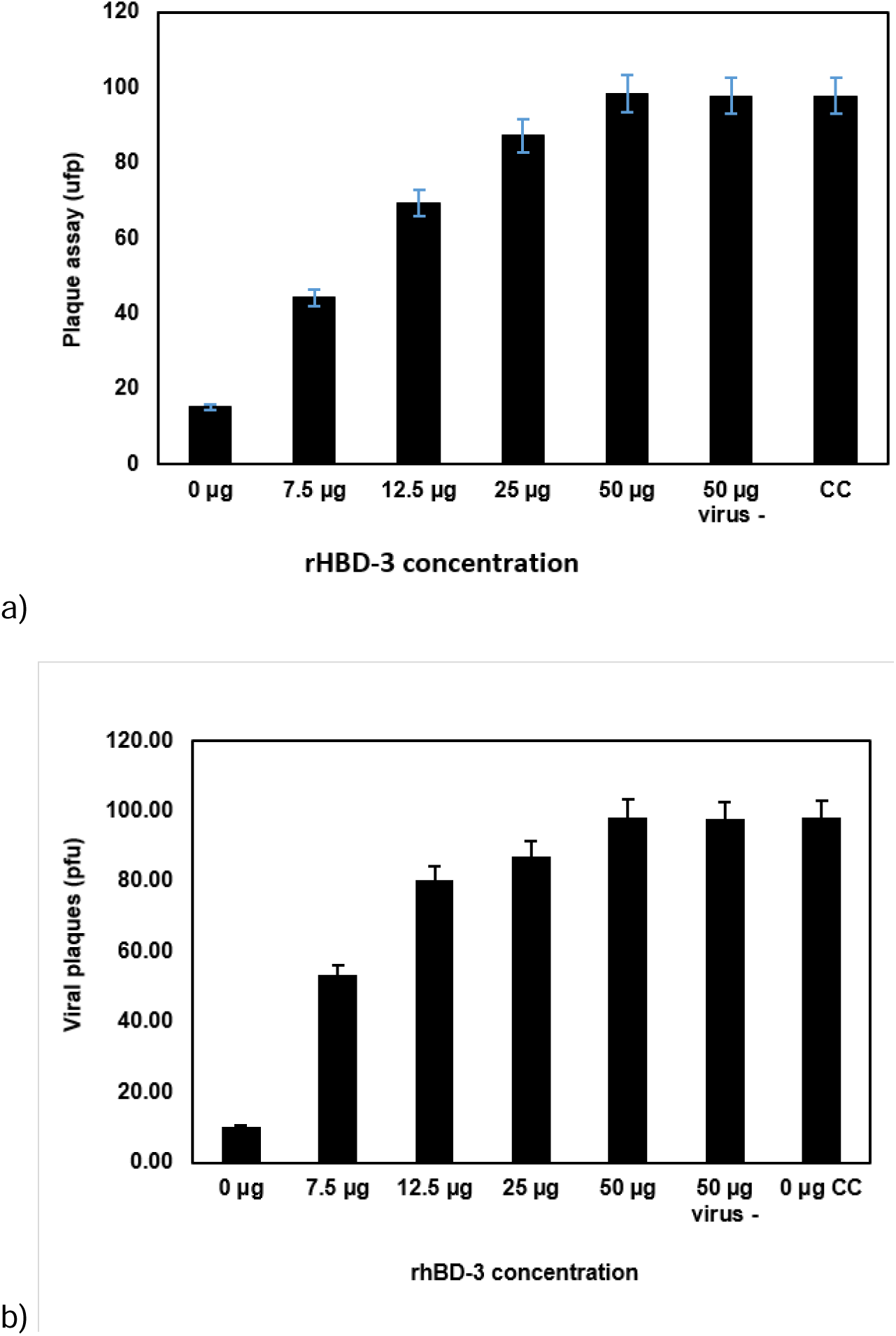
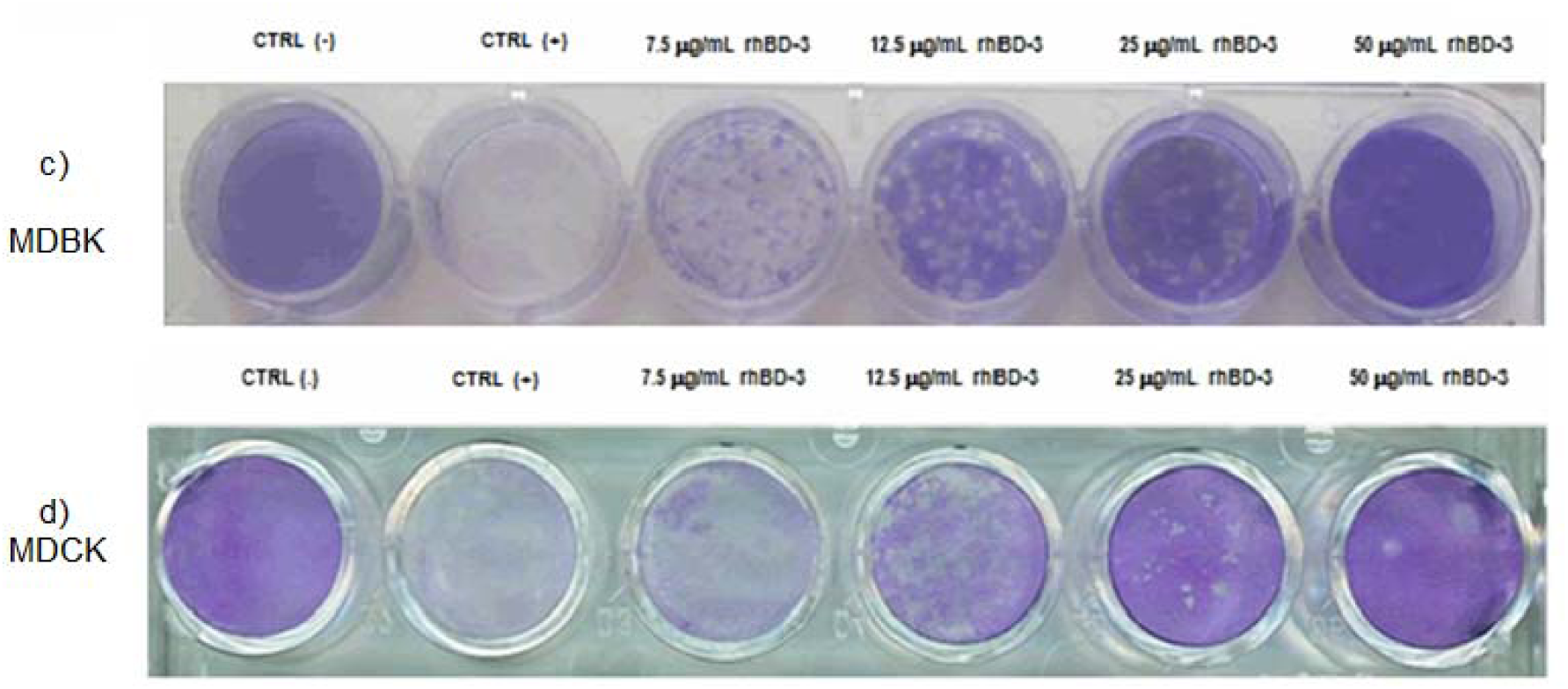
*In vitro* antiviral activity of hBD-3 against influenza A virus (IAV) in MDBK and MDCK cells. (a) Cell viability of MDBK cells treated with rhBD-3 at 7.5, 12,5, 25, and 50 μg/mL, and infected with IAV. rhBD-3 inhibited viral infection in MDBK cells in a dose-dependent manner. (b) Defensin similarly inhibited IAV in MDCK cells. Wells were run with only defensin (50 µg virus-) to assess cytotoxicity as a mechanism of action. On the graphs: CC, control cells without defensin or virus. Data correspond to the mean of three individual experiments, shown as means ± SE. c) and d) Monolayers of MDBK and MDCK cells treated with hBD-3 as indicated and infected with IAV (100 pfu). First wells are control cells not infected; in the second through sixth wells, distinct concentrations of rhBD-3 are indicated.

## Discussion

In our in vitro experiments, IAV infection significantly upregulates hBD-3 mRNA and protein expression in the human airway epithelial cell lines A549 and HEp-2. AdV infection also induces mRNA, although to a lesser degree. Our experiments further demonstrated that hBD-3, in a recombinant form, can inhibit IAV and AdV infections in A549 and HEp-2 cells, and additionally, IAV infection in epithelial cells of canine (MDCK) and bovine (MDBK) origin. Induction of hBD-3 mRNA by respiratory viruses in A549 and HEp-2 cells is a novel observation that complements previous findings indicating that respiratory virus (IAV, and rhinovirus) induce hBD-3 expression in human and murine airway epithelia [Chong *et al*., 2008; Duits *et al*., 2003; Chen *et al*., 2018]. This is the first report to show induction of hBD-3 in HEp-2 cells by IAV or AdV-5. HEp-2 cells and A549 are immortalized cell lines derived from transformed cells, and their responses to viral infection may differ from those of normal primary epithelial cells. However, these cells were chosen for experimentation for two reasons: a) they are respiratory epithelial cells of human origin, so representing the first line of defense in the respiratory system; and b) IAV and AdV replicate well in these cells. In this study, these epithelial cells yielded reproducible results, supporting the assessment of how viral infection influences hBD-3 mRNA expression and hBD-3’s antiviral activity *in vitro*.

Western blot and immunofluorescence also showed significant hBD-3 protein production during IAV infection, but not during AdV-5 infection. This could be due to the higher sensitivity of RT-PCR compared to immunofluorescence or WB, or, more specifically, to a decreased protein processing despite demonstrable mRNA production during AdV infection; the reasons for this remain to be explored. Our observations on hBD-3 production in these human cell lines during viral infection support the idea that hBD-3 may play a role in innate defense against respiratory viruses *in vivo*. Mainly because, in our model, hBD-3 mRNA was induced by two very different respiratory viruses: an enveloped virus and a naked one. Further, BDs can be induced, alone or synergistically, by pro-inflammatory cytokines such as IL-1, IL-6, and TNF-alpha, which are elevated during viral respiratory infections. Therefore, it would be interesting to investigate the interrelationships among viruses, hBD-3, and pro-inflammatory cytokines in animal models or observational studies in humans.

On the other hand, hBD-3 has shown antiviral activity against viruses from very different families: respiratory syncytial virus, herpes simplex virus, and West Nile virus [Latsko *et al*., 2022; Hazrati *et al*., 2006; Chessa *et al*., 2022]. However, there are no similar reports about IAV and AdV. The rhBD-3 used in our assays was developed in a prior study [Corrales *et al*., 2013]. This defensin was synthesized in soluble form, with over 90% purity, and demonstrated strong antibacterial activity against *Escherichia coli, Staphylococcus aureus, Pseudomonas aeruginosa*, and *Mycobacterium tuberculosis*. Additionally, this study showed *in vitro* antiviral activity against two additional viruses: a naked DNA virus, AdV, and an enveloped RNA virus, IAV, confirming the broad-spectrum antiviral properties of the hBD-3. Furthermore, defensin could be used in animal experiments, as suggested by our results with the MDBK and MDCK cells. Our observations reinforce the idea that hBD-3 is a broad-spectrum defensin exhibiting both antibacterial and antiviral properties that could play a protective role in human viral infections and could be used for treatment studies.

Regarding the mechanism of action, it is known that defensins can inhibit viruses by blocking the viral protein ligand, thereby preventing attachment and penetration steps (Gong *et al*., 2010). Defensins also exert their effects during the intracellular phases of replication [Salvatore *et al*., 2007; Smith *et al*., 2010; Quiñones *et al*., 2003; Ding *et al*., 2009]. Our *in vitro* model was designed to measure the overall effect of hBD-3. First, AdV and IAV were incubated with rhBD-3, and then the infected cells were treated with the same defensin; therefore, in our model, the two distinct actions were not separated, and both could have, wholly or partly, influenced viral infection. As a result, our model could be further used to study both aspects of the inhibitory effect. Nonetheless, immunofluorescence results showed that secreted hBD-3 and recombinant rhBD-3 migrate to the cell surface (Fig. 3i). This finding supports the idea that hBD-3 may act during the early stages of the viral replication cycle, such as during virus attachment, through steric hindrance. Taken together, our data support further studies on the role of hBD-3 in the defensive response against respiratory viruses, both in animal models and in humans.

Finally, our data open the possibility of using this molecule against respiratory viruses in future treatments, especially for emerging agents such as IAV and coronaviruses. hBD-3 could be produced *in vitro* in cell lines during post-infection assays using killed virus or by transfecting genes in *E. coli* (on a large scale, as in insulin production). On the other hand, hBD-3 could be boosted *in vivo* by pro-inflammatory cytokines during viral infection and would target the virus, helping to inhibit its replication. Further studies are needed to investigate the interrelationships between HBD-3 and the various effectors of the antiviral innate immune response that stimulate the natural production of HBD-3. Finally, hBD-3 could be combined with other antiviral agents (e.g., amantadine, ribavirin, or interferons) to explore potential synergistic effects. To advance this research, further *in vitro* and animal studies are necessary before applying these findings to humans *in vivo* clinical trials.

## Conclusions

Conclusions: Our results demonstrate that IAV and AdV-5 infections can induce the *in vitro* expression of HBD-3 in human respiratory epithelial cells, at both the mRNA and protein levels. HBD-3, in recombinant form, protected human respiratory epithelial cells against both viruses, an RNA enveloped virus and a DNA naked virus, in a dose-dependent manner. Our evidence also indicates that hBD-3 is expressed and localized on the surface of infected cells. HBD-3 also showed a species-independent inhibitory effect on IAV infection in bovine and canine epithelial cells. These findings suggest that hBD-3 plays a broad-spectrum role in defense against respiratory viruses in humans and animals and modulates the innate immune responses. Overall, this work supports the use of hBD-3 as a natural, broad-spectrum antiviral agent, either alone or combined, to treat viral infections, especially for emerging agents such as IAV and coronaviruses, for viral pandemic preparedness and response.

## Declarations

### Ethical approval

Institutional Review Board Statement not applicable.

### Informed Consent Statement

Not applicable.

### Author contributions

MAGM: Methodology, validation, investigation, data analysis and curation, original draft preparation; JS: Methodology, validation, investigation, data analysis and curation, original draft preparation; GC: Conceptualization, validation, resources, review and editing, preparation of the rhBD-3; LLCG: Methodology, validation, investigation, data analysis and curation, preparation of the rhBD-3; LMT: Conceptualization, validation, resources, review and editing. All five authors reviewed and approved the manuscript.

## Acknowledgments

We thank the Instituto de Biotecnología, Universidad Nacional Autónoma de México, and the Instituto Nacional de Enfermedades Respiratorias Ismael Cosío Villegas for the oficial primary financial support.

## Data Availability Statement

“Not applicable.”

## Availability of materials

Samples of the cells, virus, or compounds are available from the authors.

## Abbreviations

AdV-5: Adenovirus type 5
CPE: Cytopathic effect
FBS: Fetal calf serum
GSH: reduced glutathione
GSSG: oxidized glutathione
hBD-3: human beta defensin type 3
IAV: Influenza A virus
MC: Methylcellulose
MDBK: Madin-Darby bovine kidney epithelial cells
MDCK: Madin-Darby canine kidney epithelial cells
MEM: Minimum essential medium
MOI: Multiplicity of infection
pfu: Plaque-forming units
rhBD-3: recombinant human beta defensin type 3
TCID_50_: Tissue culture infectious doses 50%

